# Forebrain glucocorticoid receptor overexpression alters behavioral encoding of hippocampal CA1 pyramidal cells in mice

**DOI:** 10.1101/2022.01.13.476185

**Authors:** Swapnil Gavade, Qiang Wei, Colin Johnston, Savannah Kounelis, Klaudia Laborc, Huda Akil, Joanna L. Spencer-Segal

**Affiliations:** Michigan Neuroscience Institute, University of Michigan, Ann Arbor, MI; Department of Internal Medicine, University of Michigan Medical School, Ann Arbor, MI

## Abstract

Stress hormone signaling via the glucocorticoid receptor (GR) modulates vulnerability to stress-related disorders, but whether GR influences how the brain encodes contextual experience is unknown. Mice with lifelong GR overexpression in forebrain glutamatergic neurons (GRov) show increased sensitivity to environmental stimuli. This phenotype is developmentally programmed and associated with profound changes in hippocampal gene expression. We hypothesized that GR overexpression influences hippocampal encoding of experiences. To test our hypothesis, we performed *in vivo* microendoscopic calcium imaging of 1359 dorsal CA1 pyramidal cells in freely behaving male and female WT and GRov mice during exploration of a novel open field. We compared calcium amplitude and event rate as well as sensitivity to center location and mobility between genotypes. GRov neurons exhibited higher average calcium activity than WT neurons in the novel open field. While most neurons showed sensitivity to center location and/or mobility, GRov neurons were more likely to be sensitive to center location and less likely to be sensitive to mobility, as compared to WT neurons. More than one-third of behavior-selective GRov neurons were uniquely sensitive to location without mobility sensitivity; these uniquely center-sensitive neurons were rare in WT. We conclude that dorsal CA1 pyramidal cells in GRov mice show increased activity in a novel environment and preferentially encode emotionally salient behavior. This heightened sensitivity to a novel environment and preferential encoding of emotionally salient elements of experience could underlie differential stress vulnerability in humans with increased glucocorticoid sensitivity.

**Significance Statement (120 words maximum):** Endogenous stress hormones, glucocorticoids, are known to alter vulnerability to stress-related disorders. Here, we find that increased sensitivity to glucocorticoid via lifelong overexpression of glucocorticoid receptor in forebrain neurons (GRov) alters the encoding of environmental experiences in the hippocampus. GRov mice showed increased activity of dorsal CA1 hippocampal pyramidal cells during exploration of a novel environment and more sensitivity to emotionally relevant behaviors. These changes in how hippocampal neurons encode environmental experiences could underlie the differences in stress vulnerability in humans with genetic and epigenetic differences in glucocorticoid receptor signaling.

## Introduction

Individual differences in stress vulnerability may result from differences in neural encoding of environmental experiences. Developmental changes in stress hormone signaling via the glucocorticoid receptor (GR) alter behavioral sensitivity to the environment and contribute important vulnerability to stress-related disorders. For example, single nucleotide polymorphisms in the GR gene with evidence for increased or decreased glucocorticoid sensitivity alter vulnerability to mood disorders, substance dependence, and eating disorders (Koper et al., 2014). Similarly, methylation of the GR gene promoter, associated with early life adversity, influences GR expression, and it is associated with vulnerability to psychopathology including post-traumatic stress disorder (Palma-Gudiel et al., 2015; Yehuda et al., 2015; Kang et al., 2018). Whether developmental GR signaling influences how the brain encodes experience is unknown.

Glucocorticoid receptor is widely expressed in the brain, and mice with altered GR expression in forebrain have been developed to improve our understanding of how neuronal glucocorticoid signaling alters vulnerability to stress-related disorders (Wei et al., 2004; Boyle et al., 2005). Mice overexpressing glucocorticoid receptor in forebrain glutamatergic neurons (GRov) show enhanced behavioral sensitivity to environmental stimuli that is developmentally programmed (Wei et al., 2004, 2012; Hebda-Bauer et al., 2010). GRov mice show profound lifelong changes in gene expression in the dorsal hippocampus, a limbic structure responsible for encoding spatial, contextual, and episodic information (Wei et al., 2012). We hypothesized that developmental GR overexpression influences the encoding of experiences in the dorsal hippocampus, which may in part underlie the differential behavioral sensitivity to environmental stimuli seen in GRov mice.

CA1 pyramidal cells are the final relay of the trisynaptic circuit conveying information through the hippocampus. They encode information about the environment and behavior including context, location, and speed (Hainmueller and Bartos, 2018; Jimenez et al., 2018; Iwase et al., 2020). To determine whether GR overexpression influences encoding of environmental exploration in dorsal CA1 pyramidal cells, we used in vivo calcium imaging in freely behaving mice to measure the activity of 1359 neurons from WT and GRov mice during exploration of a novel open field. We compared hippocampal CA1 pyramidal cell calcium activity and behavioral sensitivity to mobility and location between genotypes.

## Methods

### Subjects

GRov mice and WT littermates were generated and maintained as previously described (Wei et al., 2012). Mice were maintained on a 14:10 light/dark cycle with imaging sessions performed during the light phase. Procedures were approved by the University of Michigan Institutional Animal Care and Use Committee and were conducted in accordance with the NIH Guide for the Care and Use of Laboratory Animals (2011).

### In vivo microendoscopic calcium imaging

In vivo calcium imaging in freely behaving mice was performed using the nVista 2.0 miniature endoscope (Inscopix, Inc.) following the procedures described in detail elsewhere (Resendez et al., 2016). Young adult male and female GRov and WT mice underwent injection of AAV5-CamKIIa-GCamp6f (Addgene) (300 nL diluted 1:5 in artificial CSF AP −2.05, ML 1.75 from bregma; DZ −1.3 from skull) and implantation of a 4 x 1 mm GRIN lens (at AP −1.95, ML 1.6 from bregma, DZ −1.55 from skull) over the dorsal hippocampal CA1 pyramidal cell layer (**Figure 1A-B**).

**Figure 1.**
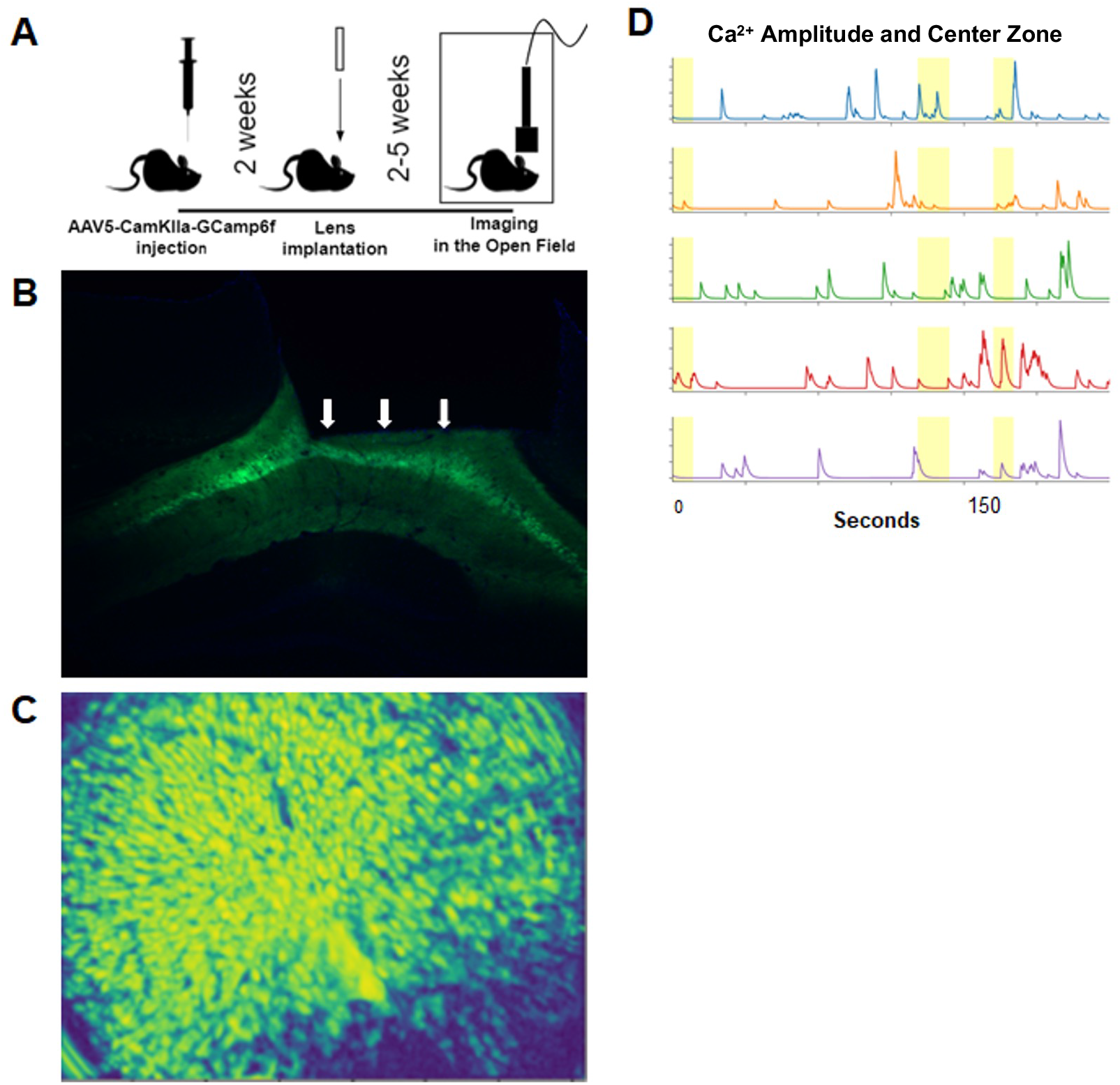
**A**, Schematic of experiment. **B**, Histology demonstrating GCamp6f expression in dorsal CA1 pyramidal cells. White arrows delineate the lower lens border. **C**, Representative recorded frame. **D**, Example traces from 5 cells in mouse during open field exploration. Yellow areas show when the mouse was in the center of the open field.

For recordings, mice were allowed to explore a large (72 cm x 72 cm) brightly lit (200 lux) open field for 10 minutes. Behavior was recorded and synchronized with calcium imaging using Noldus Ethovision XT (Version 12) (**Figure 1C**). At least 3 days after the completion of behavioral testing, mice were perfused with PBS followed by 4% paraformaldehyde and brains were postfixed in paraformaldehyde for 24 hours prior to removal of the lens. Histology was used to verify GCamp6f expression in dorsal CA1 pyramidal cells and accurate placement of the lens over the dorsal CA1 region.

### Data processing

Downsampling (by 2 in space and time), cropping and trimming of videos were performed using Inscopix Data Processing Software (IDPS 1.6.0). CaImAn, an open-source tool for scalable calcium imaging data analysis (Zhou et al., 2018), was used to perform background subtraction, cell identification and trace extraction on collected calcium imaging data using the Python toolbox. Manual curation of the identified cells was performed to ensure that the final accepted cells were unique and exhibited typical calcium dynamics. Event detection through deconvolution was performed using the Online Active Set method to Infer Spikes method (Friedrich et al., 2017).

To determine the behavioral sensitivity of hippocampal neurons, calcium activity was aligned with behavior including location (center and periphery) and mobility, which was assigned bins of high or low mobility (velocity >= 5 cm/s) and < 5 cm/s, respectively. Calcium amplitude was shuffled for each animal over behavior bins to generate a shuffled data distribution, a “null” distribution where the calcium activity will not be sensitive to any behavior. For each neuron, a ratio was calculated for the average calcium amplitude in the center/periphery, and high/low mobility. Thresholds for behavioral sensitivity were defined as cells with a ratio lower than the 1st percentile or higher than the 99th percentile, based on the shuffled data distribution within genotype (**Figure 2A**). These cells were considered mobility or location selective and represent cells that selectively increase or decrease their calcium activity when the mouse increases speed or enters the center of the open field (positive and negative speed cells, and center and periphery selective cells). Whether calcium activity anticipated a behavioral change was examined by examining calcium activity during the one second period before the mouse entered the center zone relative to the average calcium activity in the periphery zone.

**Figure 2.**
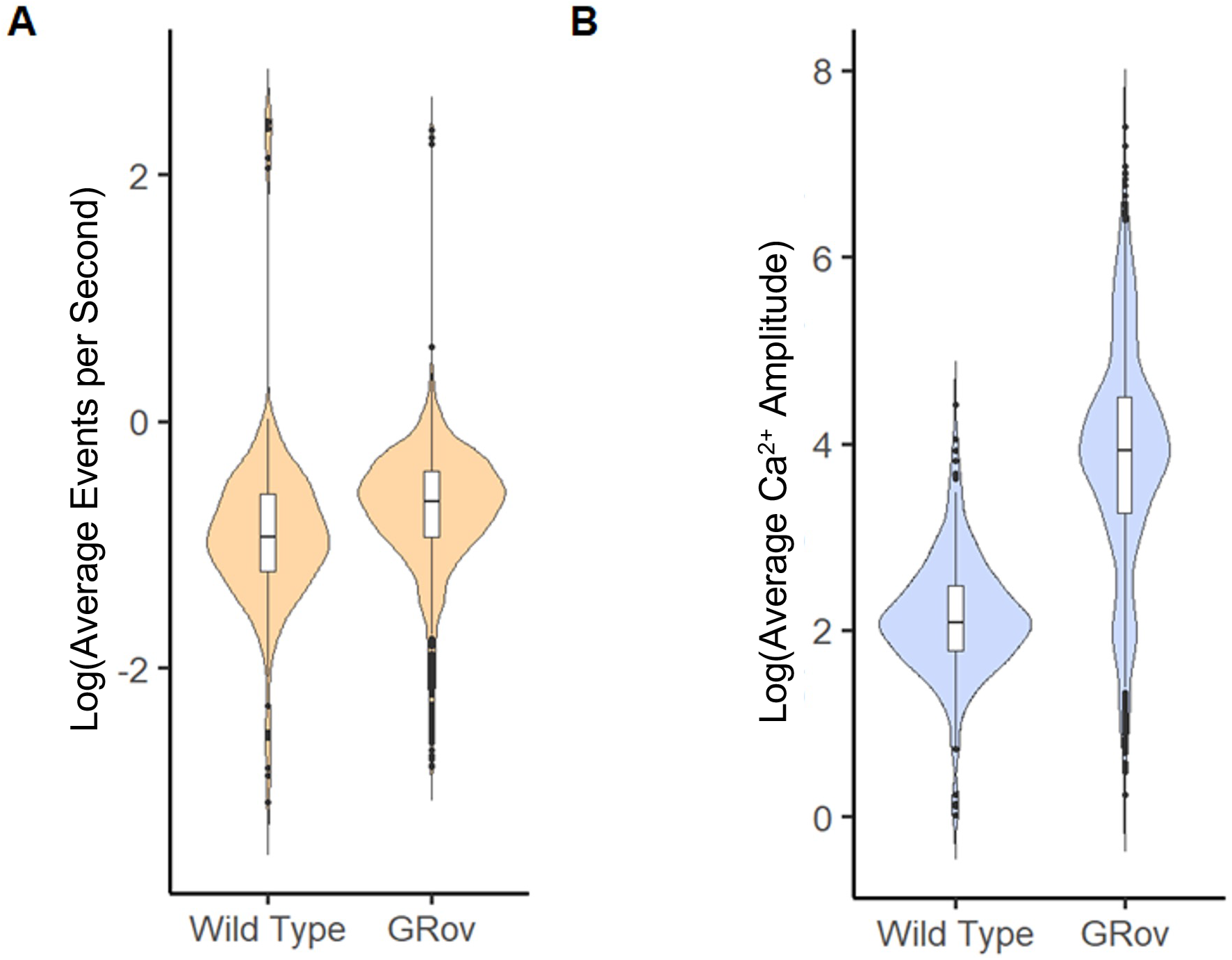
**A,** Example distributions of the ratio of calcium event rate in high/low mobility behavior bins for each neuron in original and shuffled data. Vertical dotted lines show thresholds for 1% and 99% of the distribution based on the shuffled data within genotype. **B-C**, Example traces of one center selective neuron and one positive speed neuron. Blue trace shows animal velocity, while the highlighted yellow region shows when the animal entered the center.

### Experimental Design and Statistical Analyses

8 mice (2 WT male, 2 WT female, 3 GRov male, and 1 GRov female) exhibited fluorescence with dynamic activity and identifiable cells, and all of these mice were included in the analysis. In total, activity from 260 neurons were identified for WT and 1099 for GRov.

To compare overall calcium activity between WT and GRov neurons, log transformed calcium amplitude and event rate were compared using linear regression models in R studio (1.4.1717) where the dependent variable was cell activity and independent variable was genotype (**Table 1**).

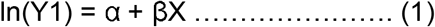

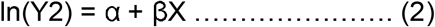

**Table 1.** Output of linear regression models used to compare average calcium amplitude and event rate for individual WT and GRov neurons in the novel open field.

| **(A) Model 1 : log(Y1) ~ genotype** |  |  |  |
| --- | --- | --- | --- |
| Coefficients: |  |  |  |
|  | Estimates | Std. error | Pr(>\|t\|) |
| (Intercept) | 2.9942 | 0.03921 | <0.0000000000000002 *** |
| Genotype WildType - Grov | -0.8566 | 0.03921 | <0.0000000000000002 *** |
| **(B) Model 2 : log(Y2) ~ genotype** |  |  |  |
| Coefficients: |  |  |  |
|  | Estimates | Std. error | Pr(>\|t\|) |
| (Intercept) | -0.80653 | 0.01943 | < 0.0000000000000002 *** |
| Genotype WildType - Grov | -0.08829 | 0.01943 | 0.00000602 *** |

Where,

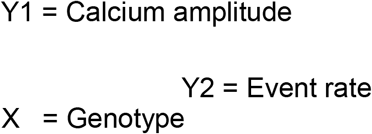

The behavioral sensitivity between GRov and wild type cells was evaluated by comparing the fraction of CA1 neurons in each genotype showing sensitivity to either speed or location in the open field using Fisher’s exact test.

Statistical analyses were performed using Python. Figures were created by Python and R and formatted in Adobe photoshop (22.5.1).

## Results

We recorded dorsal CA1 pyramidal cell calcium activity from 4 WT and 4 GRov mice during exploration of a novel open field to understand differences in overall cellular activity in a novel environment, as well as behavior sensitivity (**Figure 1**). First, to determine whether overall calcium activity differed between genotypes, we examined average calcium amplitude and event rate between WT and GRov neurons. During exploration of a novel open field, GRov neurons exhibited higher average calcium amplitude and event rate than WT neurons (**Figure 2A-B** and **Table 1**), demonstrating increased activity in dorsal CA1 pyramidal cells in GRov compared to WT during novel open field exploration.

We used behavior sensitivity analysis to identify cells that showed sensitivity to mobility or center location in the open field, sensitivities expected in dorsal CA1 neurons based on previous observations (Jimenez et al., 2018; Iwase et al., 2020) (**Figure 2**). Cells that preferentially increased their calcium activity when the animal was moving faster or slower were termed “high mobility cells” or “low mobility cells,” while cells that increased their activity in the center or periphery were termed “center cells” or “periphery cells,” respectively. Overall, the vast majority of dorsal CA1 neurons were sensitive to mobility and/or location: 90.60% in GRov and 95.33% in WT.

Consistent with prior observations, the majority of dorsal CA1 pyramidal cells were sensitive to mobility in the open field, and the same was true for location (**Figure 3**). More cells met criteria for behavior sensitivity using the calcium amplitude than the event rate measure; further inspection of the data demonstrated that all the behavior-sensitive cells identified in the event rate analysis were also identified as behavior-sensitive cells in the calcium amplitude analysis, confirming our suspicion that, rather than returning different sets of cells, these two analyses capture similar information about behavior sensitivity with event rate being a more stringent measure.

**Figure 3.**
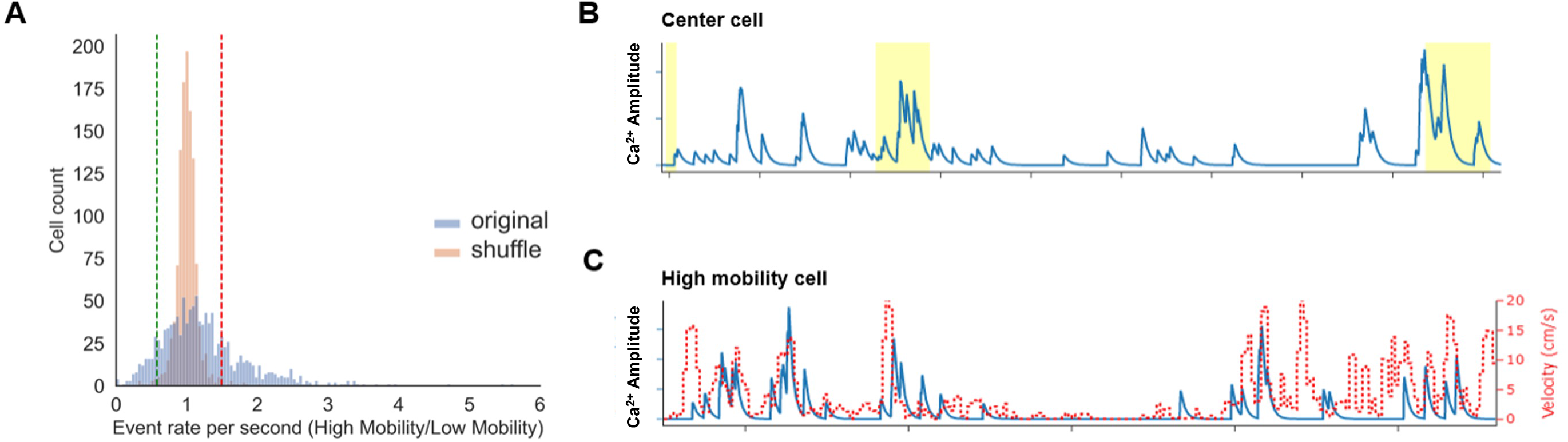
Bar graph showing the location and mobility of sensitive cells based on calcium amplitude and event rate..

Despite the high number of behavior-sensitive cells in each genotype, the proportion of mobility- and location-sensitive neurons was different between genotypes (**Figure 4**). More WT than GRov neurons displayed mobility sensitivity: 88% of WT, 63% of GRov neurons based on calcium amplitude (P=0.001), and 59% of WT, 53% of GRov neurons based on event rate (P=0.4542). The difference in mobility sensitivity in calcium amplitude appeared driven by a decrease in low mobility cells in GRov: 40% of the neurons were low mobility cells in WT, compared with 21% in GRov (P<0.001). On the other hand, more GRov than WT neurons displayed center/periphery location sensitivity: 68% of WT/79% of GRov neurons displayed location sensitivity based on calcium amplitude (P=0.18), and 31% of WT, 54% of GRov neurons displayed location sensitivity based on calcium event rate (P=0.000017). The difference in location sensitivity in event rate appeared driven by an increase in periphery cells in GRov: 21% of GRov and 0% of WT neurons were periphery cells (P<0.001). In summary, dorsal CA1 pyramidal cells in GRov showed more location sensitivity, and less mobility sensitivity, than WT neurons in the novel open field.

**Figure 4.**
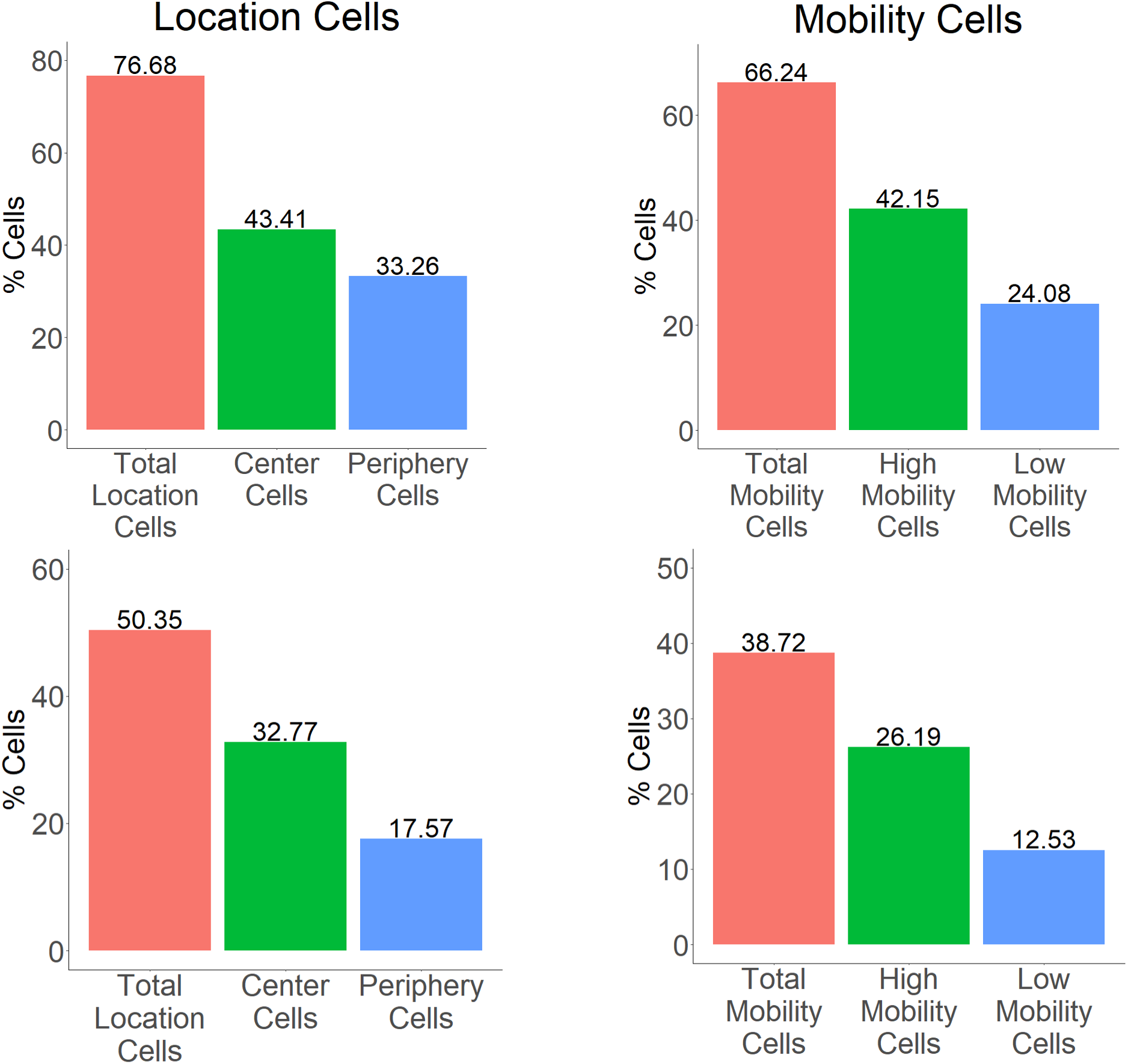
Graph shows the percentage of mobility- and location-sensitive neurons based on calcium amplitude, and event rate *P<0.05 for GRov and WT.

Next, we looked at shared sensitivities between mobility and location to determine the overlap in behavioral sensitivities within individual neurons. Individual CA1 pyramidal cells frequently encoded both speed and location (**Figure 5**). For both genotypes, center cells were more likely to also be high mobility cells than low mobility cells; the opposite was true for periphery cells, which were more often low mobility cells than high mobility cells **(Figure 6)**. Since mice generally move at higher speeds through the center of the open field, this raised the possibility that the speed sensitivity analysis could be confounded by location. When the analysis was repeated excluding center times, it did not substantially change individual neuron categorization, suggesting a true shared behavioral sensitivity rather than confounding.

**Figure 5.**
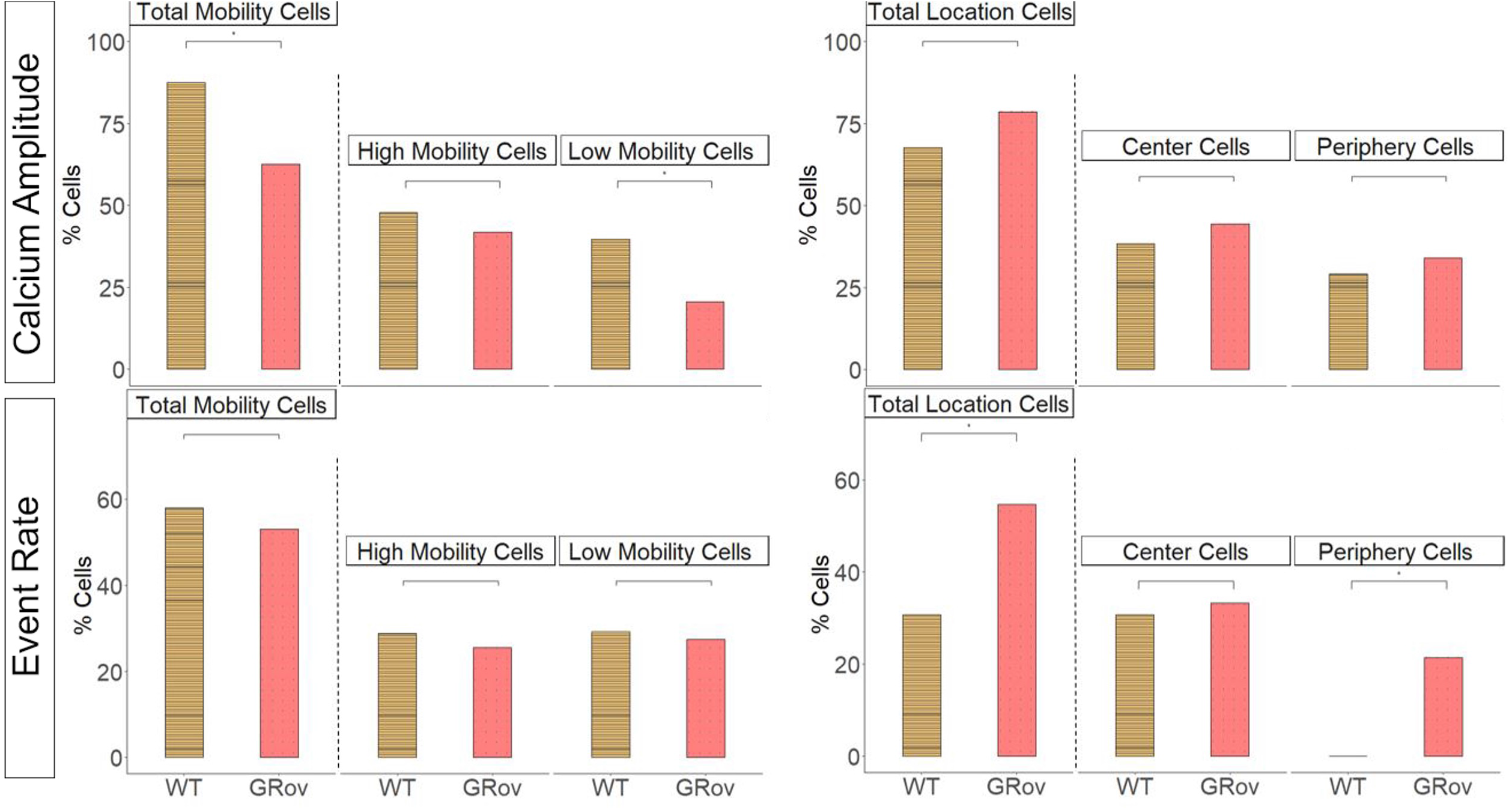
Venn diagrams show the fraction of overlapping neurons between speed and location sensitive neurons. + represents high mobility cells, - represents negative speed neurons, CS= center cells, PS = periphery cells)

**Figure 6.**
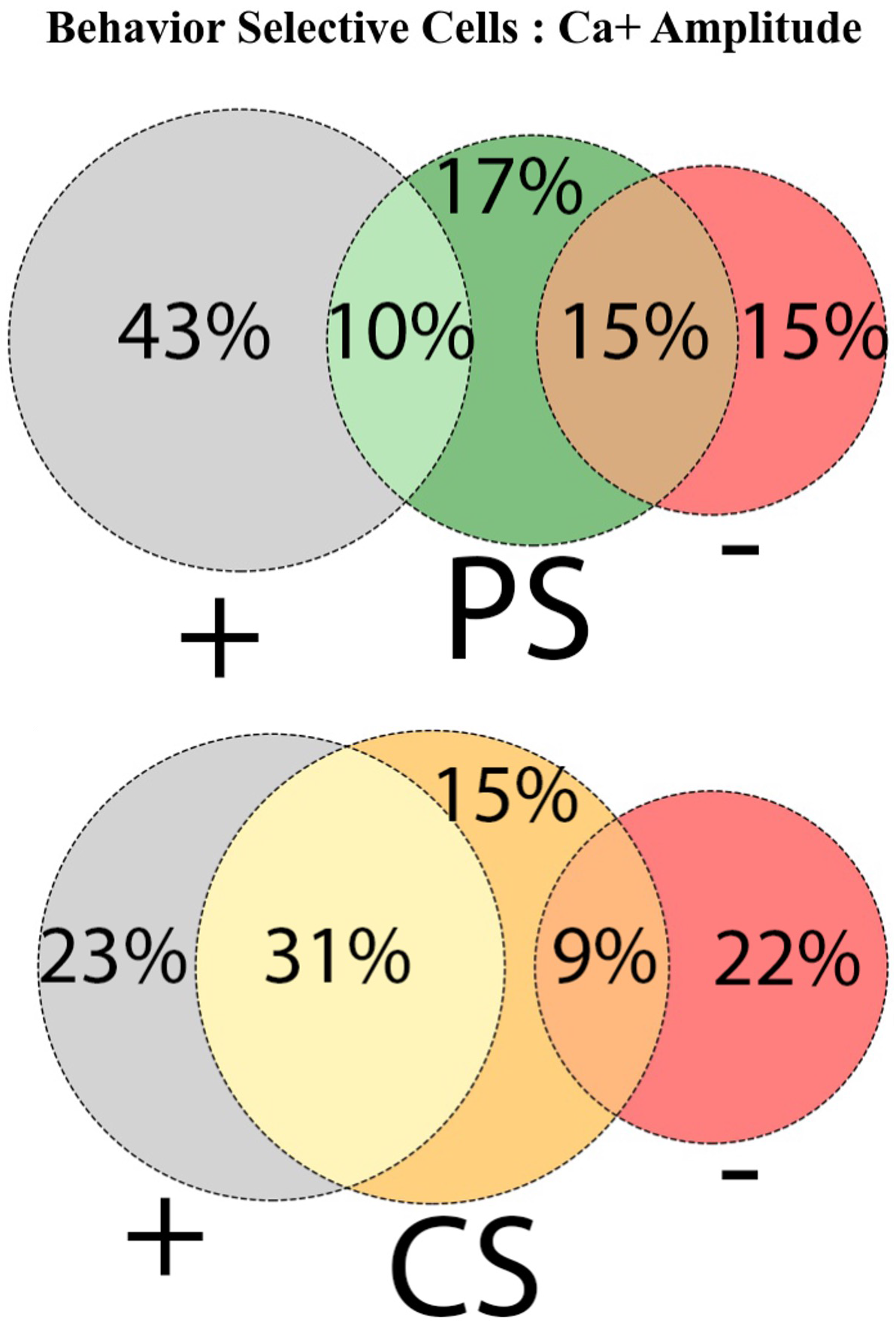
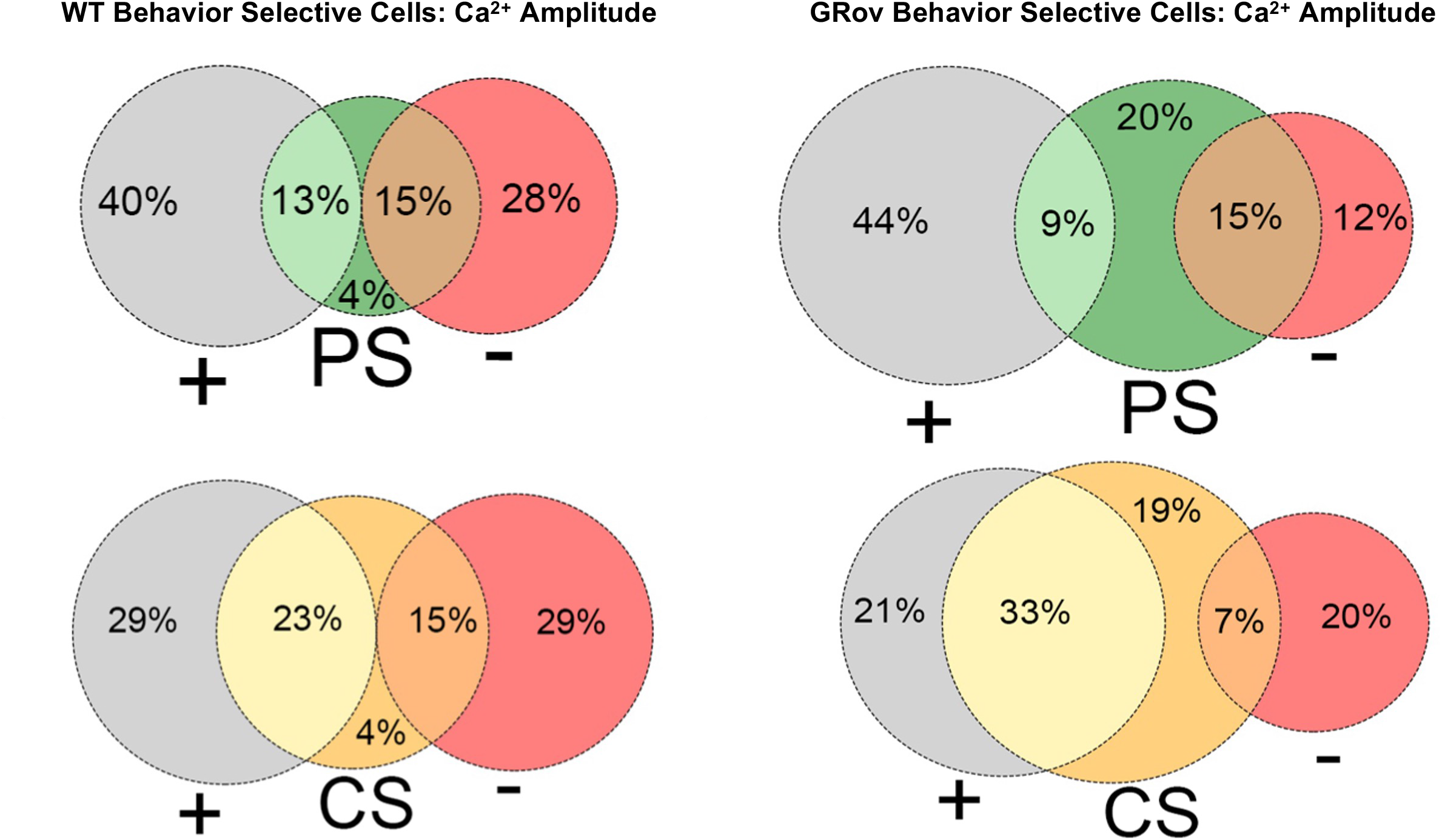
Venn diagrams for each genotype show the fraction of overlapping neurons between speed and location sensitive neurons.

The degree of overlap in behavior sensitivities was different between genotypes (**Figure 6**). Almost all location cells in WT were also mobility cells, with only 4% uniquely selective for either periphery or center. In contrast, more GRov neurons were uniquely selective for the periphery (20%) or center (19%) without displaying any mobility sensitivity. To further understand the distribution of cells with unique behavior sensitivity to either mobility or location between genotypes, we compared the proportion of all identified neurons with unique mobility or location sensitivities between genotypes (**Figure 7**). More WT than GRov neurons showed unique mobility sensitivity: high mobility cells comprised 15% of all neurons in WT and 8% in GRov (P=0.004), while low mobility cells comprised 12% of all neurons in WT and 4% in GRov (P<0.001). In contrast, more GRov than WT neurons showed unique location sensitivity: center cells comprised 4% of all neurons in WT and 15% in GRov (P<0.001), while periphery cells comprised 4% of all neurons in WT and 14% in GRov (P<0.001). In other words, about 1 in 3 of all detected GRov neurons showed unique sensitivity to center/periphery location without any mobility sensitivity, while these unique location-sensitive cells were rare (less than 1 in 10) in WT neurons.

**Figure 7:**
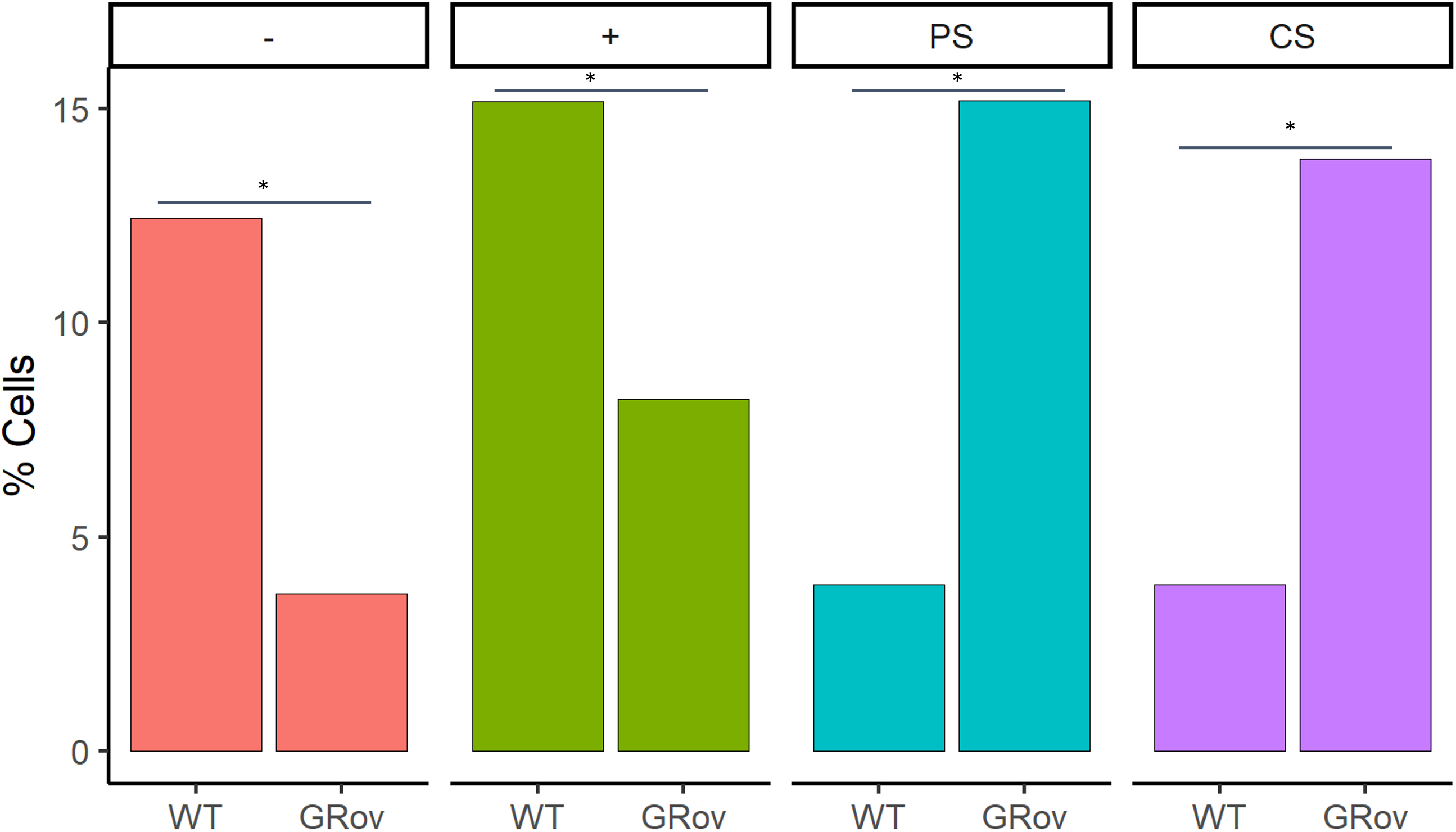
Fraction of uniquely identified low mobility (-), high mobility (+), center and periphery cells in WT and GRov as a percentage of the total accepted cells. *P<0.01

Finally, we sought to understand whether dorsal CA1 neurons primarily represent behavior and experience, or whether they anticipate behavior and thus may plausibly drive it. To determine whether CA1 neurons might drive center exploration, we analyzed calcium activity during the 1 second before the animal entered the center and compared activity in the pre-center and periphery bins in center cells (**Figure 8**). The majority (69.84%) of center cells showed higher calcium activity during pre-center bin in comparison to calcium activity in the overall periphery zone. This anticipation suggests that many of these cells could drive center exploration. This percentage was similar between genotypes (65% in WT, 71% in GRov).

**Figure 8.**
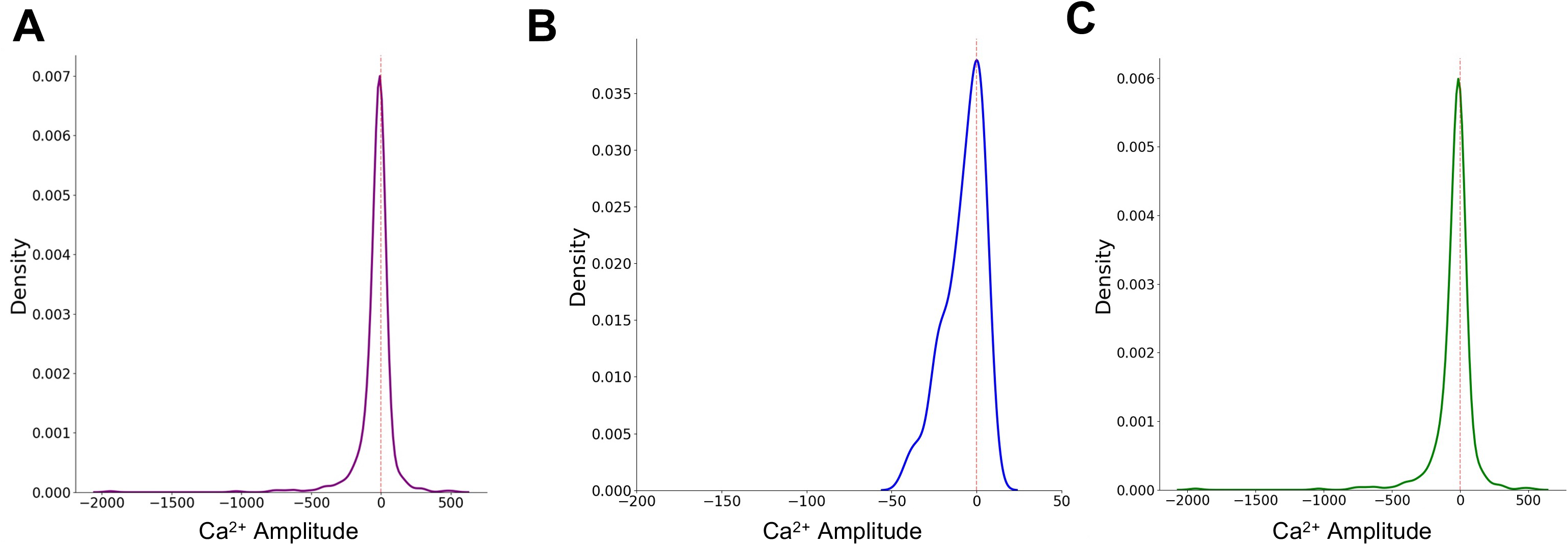
**A**, Distribution of the change in average calcium activity in pre-center bin from average calcium activity in rest of the periphery bin. **B-C**; Distribution of change in average calcium activity in WT (**B**) and GRov (**C**).

## Discussion

Here we demonstrate that lifelong overexpression of GR in forebrain glutamatergic neurons alters behavioral encoding in dorsal CA1 pyramidal cells during novel open field exploration. Compared to WT, GRov neurons displayed overall higher calcium activity and preferential sensitivity to center/periphery location over mobility.

The finding of an overall increase in the calcium activity in CA1 neurons during novel open field exploration most likely represents an increase in firing rate, given the correlation between calcium dynamics and cell electrical activity (Chen et al., 2013). However, it could also reflect differences in calcium handling between the two genotypes (Wei et al., 2012). It does not reveal whether this increased activity is a basal property of increased excitability in GRov neurons, or whether it reflects a heightened response to the novel environment, as CA1 neurons are known to increase their activity in a novel environment (Karlsson and Frank, 2008). The increased CA1 pyramidal cell activity during exploration of a novel environment in GRov recalls the increased behavioral sensitivity to the environment, including decreased overall center exploration, previously seen in GRov mice (Wei et al., 2004, 2012; Hebda-Bauer et al., 2010). While GRov mice had more center and periphery cells than WT, the relative increase in periphery cells in GRov could explain decreased center exploration, if these cells do indeed drive behavior. Indeed, our analysis demonstrated anticipation of center exploration in most center neurons, suggesting that at least some location-sensitive neurons may drive behavior.

The findings of high rates of mobility and location selectivity of CA1 pyramidal cells are consistent with previous reports (Jimenez et al., 2018; Iwase et al., 2020). Our findings are novel in their demonstration of high rates of shared mobility and center sensitivities in CA1 pyramidal cells, which have not previously been compared in the same neuronal population. Between genotypes, GRov neurons displayed more location sensitivity, and less mobility sensitivity, than WT neurons. While center and periphery location are often interpreted in the context of their relevance to innate anxiety states, it was previously suggested that the location sensitivity of dorsal CA1 cells represents spatial, and not emotional, information (Jimenez et al., 2018). Intriguingly, we found a high proportion of neurons that were uniquely location sensitive in GRov, with no mobility sensitivity. This raises the possibility of differential tuning of the dorsal hippocampus in GRov toward more emotionally salient stimuli, a hypothesis which will need to be tested.

In conclusion, lifelong overexpression of GR in forebrain neurons alters the information encoded by CA1 pyramidal cells. They display higher calcium activity in a novel environment and preferential encoding of the emotionally relevant center/peripheral location in a novel open field. We suggest that this increased activity and differential encoding of experience in a novel context by dorsal CA1 pyramidal cells contributes to the differential behavioral sensitivity and more emotionally labile phenotype in GRov mice. The findings suggest that humans with developmental differences in glucocorticoid receptor signaling may also demonstrate differential encoding of experiences within the hippocampus, contributing to differential vulnerability to stress-related disorders. In the future, we recommend more consideration of the contribution of the dorsal hippocampus and its innate biases in episodic and contextual encoding in stress vulnerability.

## Acknowledgements

The authors would like to acknowledge Mark Reimers, PhD, René Hen, PhD, and Jessica Jimenez, MD, PhD for their technical expertise and advice.

## Conflicts of Interest

The authors declare no competing financial interests.

## Funding Sources

Hope for Depression Research Foundation (H.A.), NIH grant MH116267 (J.S.S.), Brain and Behavior Research Foundation (J.S.S.)

## Notes

### Competing Interest Statement

The authors have declared no competing interest.

### Summary of Updates

Author order

